# HY5 and COP1 function antagonistically in the light-dependent regulation of nicotine biosynthesis in tobacco

**DOI:** 10.1101/2024.02.28.582661

**Authors:** Deeksha Singh, Shambhavi Dwivedi, Nivedita Singh, Prabodh Kumar Trivedi

## Abstract

Nicotine constitutes approximately 90% of the total alkaloid content within the *Nicotiana* species, rendering it the most prevalent alkaloid. While the majority of genes responsible for nicotine biosynthesis express in root tissue, the influence of light on this process through shoot-to-root mobile ELONGATED HYPOCOTYL 5 (HY5) has been recognized. CONSTITUTIVE PHOTOMORPHOGENIC1 (COP1), a key regulator of light-associated responses, known for its role in modulating HY5 accumulation, remains largely unexplored in its relationship to light-dependent nicotine accumulation. Here, we identified NtCOP1, a COP1 homolog in *Nicotiana tabacum*, and demonstrated its ability to complement the *cop1* mutant in *Arabidopsis thaliana* at molecular, morphological, and biochemical levels. Through the development of NtCOP1 overexpression (NtCOP1OX) plants, we observed a significant reduction in nicotine and flavonol content, inversely correlated with the down-regulation of nicotine and phenylpropanoid pathway. Conversely, CRISPR/Cas9-based knockout mutant plants (*NtCOP1^CR^*) exhibited an increase in nicotine levels. Further investigations, including yeast-two hybrid assays, grafting experiments, and western blot analyses, revealed that NtCOP1 modulates nicotine biosynthesis by targeting NtHY5, thereby impeding its transport from shoot-to-root. We conclude that the interplay between HY5 and COP1 functions antagonistically in the light-dependent regulation of nicotine biosynthesis in tobacco.

**Highlight:** Characterization of CONSTITUTIVE PHOTOMORPHOGENIC1 (COP1) overexpressing and CRISPR/Cas9-based mutant plants suggests the intricate role of COP1 in modulating nicotine biosynthesis in tobacco.

## INTRODUCTION

Nicotine, the primary alkaloid found in Nicotiana tabacum, constitutes a substantial 90% of the total alkaloids present (Dewey and Xie, 2013). Synthesized initially within root cortical cells, nicotine undergoes a series of enzymatic transformations before being stored in vacuoles (Shoji *et al*., 2009). Its moveent continues as it is transported to leaf vacuoles through the xylem, where it acts as a potent defense mechanism against herbivores (Steppuhn *et al*., 2004). Structurally, nicotine is composed of a pyridine and a pyrrolidine ring (Hibi *et al*., 1994). Several enzymes, including aspartate oxidase and quinolinate synthase (QS) (Katoh *et al*., 2006), as well as quinolinate phosphoribosyl transferase (QPT) (Sinclair *et al*., 2000), catalyze the formation of the pyridine ring from nicotinic acid (Hashimoto and Yamada, 1994). The pyrrolidine ring, on the other hand, is derived either directly from putrescine via ornithine decarboxylase (ODC) (Imanshi *et al*., 1998) or indirectly from arginine through arginine decarboxylase (ADC) (Chattopadhyay *et al*., 1998). Putrescine N-methyltransferase (PMT) facilitates the conversion of putrescine into N-methyl-putrescine (Riechers *et al*., 1999), which is then oxidized by N-methyl-putrescine oxidase (MPO) and cyclized to form the N-methyl pyrrolinium cation (Naconsie *et al*., 2014). Through a complex series of reactions involving NADPH-dependent PIP reductase A622 (De Boer, 2009) and berberine bridging enzyme-like (BBL) proteins, this cation combines with nicotinic acid to produce nicotine (Kajikawa *et al*., 2011).

Previous studies on the regulation of nicotine biosynthesis have primarily focused on two aspects: structural genes within the nicotine pathway and hormonal regulation. Jasmonic Acid (JA) has been identified as a key player in stimulating nicotine biosynthetic genes by promoting the proteasomal degradation of JA-ZIM domain (JAZ) repressor proteins (Shoji *et al*., 2008). These repressors, including NtMYC2a/2b (Zhang *et al*., 2012) and R2R3-MYB proteins such as AtMYB21/24/305a (Song *et al*., 2011; Bian *et al*., 2022), act to inhibit basic helix–loop–helix transcription factors in Nicotiana. Investigations into the PMT1a promoter have underscored the significance of the GAG region, comprising a G-box, an AT-rich motif, and a GCC-box-like element essential for JA induction (Xu *et al*., 2004). While BHLH and MYB factors bind to the G-box (Shoji and Hashimoto, 2011a), NtERFs like NtERF32 (Sears *et al*., 2014), NtERF189 (Shoji *et al*., 2010), and NtERF221 (Liu *et al*., 2019) interact with the GCC box-like element. Recent studies have identified NtMYB305a as a regulator binding to the AT-rich motif, exerting a positive effect on nicotine biosynthesis (Bian *et al*., 2022). In a previous investigation, we explored the role of the light signaling component NtHY5 in nicotine biosynthesis. Our findings unveiled the positive regulatory function of NtHY5, as mutations in HY5 resulted in a significant decrease in nicotine biosynthesis and accumulation (Singh *et al*., 2023). However, the involvement of other light-associated networks has not been studied as yet.

In the absence of light, plants undergo a transition from photomorphogenesis to skotomorphogenesis, a process orchestrated by the central repressor CONSTITUTIVE PHOTOMORPHOGENIC1 (COP1)/DE ETIOLATED/FUSCA proteins (Yi and Deng, 2005). COP1, functioning as a RING E3 ligase, translocates into the nucleus under dark conditions, where it targets key light signaling components such as ELONGATED HYPOCOTYL 5 (HY5), HYH, LONG HYPOCOTYL IN FAR-RED 1 (HFR1), and LONG AFTER FAR-RED LIGHT 1 (LAF1) for degradation via the 26S proteasome pathway. Downstream of COP1, transcription factors HY5, HYH, and PHYTOCHROME-INTERACTING FACTOR (PIF) coordinate two distinct signaling branches in the photomorphogenic response of *Arabidopsis thaliana* (Castillon *et al*., 2007; Lau and Deng, 2012). Members of the bZIP transcription factor family, HY5, and HYH, regulate the expression of numerous genes associated with photomorphogenesis (Osterlund *et al*., 2000; Holm *et al*., 2002; Lee *et al*., 2007; Zhang *et al*., 2011). In the dark, COP1 relocates from the nucleus to the cytoplasm, where it accumulates to facilitate degradation through polyubiquitination, thereby promoting skotomorphogenesis (Holm *et al*., 2002; Hoecker *et al*., 2017).

Extensive research has suggested role of COP1 as a repressor across various biological processes, including photomorphogenic development, flavonoid biosynthesis, embryo development, the transition from vegetative to reproductive phases, and anthocyanin biosynthesis, spanning plant species such as *Arabidopsis thaliana, Oryza sativa, Pyrus bretschneideri,* and *Malus domestica* (Li *et al*., 2013; Maier *et al*., 2012; Tanaka *et al*., 2011; Kim *et al*., 2017; Wu *et al*., 2019; Bhatia *et al*., 2021). While our previous work elucidated the role of NtHY5 in nicotine biosynthesis (Singh *et al*., 2023), there remains a gap in understanding the involvement of COP1 in the biosynthesis of these molecules. Addressing this gap, our study investigates and unveils the role of NtCOP1 in nicotine and flavonoid biosynthesis. Through a comprehensive analysis employing various overexpressing lines, CRISPR/Cas9-based mutants, and grafting experiments, we demonstrate that HY5 and COP1 operate antagonistically in the regulation of nicotine biosynthesis in tobacco.

## MATERIALS AND METHOD

### Isolation and sequence analysis of NtCOP1

To identify the ortholog of AtCOP1 in tobacco, BLAST searches were performed on National Center for Biotechnology Information (http://www.ncbi.nlm.nih.gov/BLAST). The sequence was retrieved and amino acid sequence was analyzed for the N terminal ring motif, alpha helical coiled coil domain and C-terminal WD-40 domain which consist of substrate binding region. Conserved domain database (CCD) was employed to ensure the conserved domains of the identified NtCOP1 protein. The genes retrieved were checked for the presence of the requisite domains using InterProScan (Quevillon *et al*., 2005) (http://www.ebi.ac.uk/Tools/pfa/iprscan/). Amino acid sequence of COP1 proteins from different plant species which belong to different families was extracted to construct the phylogenetic tree using the Neighbor-Joining (NJ) algorithm in MEGA6.0 software (Kumar *et al*., 2008).

### Plant material, growth conditions and treatments

Leaves of *Nicotiana tabacum* cv. Petit Havana (NtPH) have been used for raising the transgenic and genome-edited plants. Tobacco plants were grown in glass house at 25°C±2°C and 16 h/8 h light-dark photoperiods for harvesting of tissues belonging to different developmental stage. For dark treatment, seeds were surface sterilized using 70% EtOH for 1 min and then dipped in 50% bleach for 4 min after that washed repeatedly with autoclaved Milli-Q water, and grown for 10 days in absence of light, 24°C temperature, and 60% relative humidity on one-half-strength Murashige and Skoog (MS) medium (Sigma) plates after stratification for 2 to 3 d at 4°C in the dark and subsequently transferred to culture room with 16 h/8 h light-dark photoperiods and 150–180 mm m^–2^s^–1^ light intensity. For light-to-dark treatment, glass house grown 5-month-old tobacco plants were completely covered with black paper for 24h before sampling. The experiment was performed by using three independent replicates. Arabidopsis (Col-0) was used as the wild-type plant for the complementation study. Seeds were surface sterilized and after stratification transferred to a growth chamber under controlled conditions of 16-h-light/8-h-dark photoperiod cycle, 22°C temperature, 150 to 180 µmol m^-2^s^-1^ light intensity, and 60% relative humidity for 10 days unless mentioned otherwise.

### Plasmid construction and generation of *Arabidopsis* and tobacco transgenic and mutant plants

To overexpress *NtCOP1*, the *NtCOP1* was amplified from the single-stranded complementary DNA (cDNA) library prepared from 10-day-old tobacco seedlings through PCR using specific primers. The full-length open reading frame of the NtCOP1 cDNA, under the control of Cauliflower Mosaic Virus 35S (CaMV35S) promoter in the binary vector pBI121 (Clontech, USA) was transferred into *Agrobacterium tumefaciens* strain GV3101 and used to transform *Arabidopsis* by floral dip method and tobacco plants by leaf disk method (Horsch *et al*., 1985). Several transgenic tobacco lines constitutively expressing NtHY5 cDNA were selected on the basis of RT-PCR. Seeds were harvested, sterilized and plated on solid half-strength MS medium supplemented with 100 mg/l kanamycin. Antibiotic resistant plants were shifted to the glass house and grown until maturity.

For generation of mutant plants of *NtCOP1*, CRISPR-Cas9 technology was used. A 20-bp gRNA was selected from the identified coding region. *NtCOP1* gRNAs was cloned into the binary vector pHSE401 using the BsaI restriction site (Xing *et al*., 2014). This vector contains Cas9 endonuclease-encoding gene under dual CaMV35S promoter as well as genes encoding neomycin phosphotransferase and hygromycin phosphotransferase as selection markers. All the constructs were sequenced from both the orientations using plasmid-specific forward and vector reverse primers after that transferred to GV3101 and then transformed to tobacco

### Gene expression analysis

Total RNA was isolated using a Spectrum Plant Total RNA kit (Sigma Aldrich) following the manufacturer’s instruction. For qRT-PCR analysis, 2 µg of DNA free RNA was reverse transcribed using a RevertAid H minus first-strand cDNA synthesis Kit (Fermentas) according to the manufacturer’s instructions. qRT-PCR was performed using Fast Syber Green mix (Applied Biosystems) in a Fast 7500 Thermal Cycler instrument (Applied Biosystems). Expression was normalized using Tubulin and analyzed through the comparative CT method (Liva and Schmittgen 2001). The oligonucleotide primers used to study expression of different genes were designed using the Primer Express 3.0.1 tool (Applied Biosystems) and information is provided in **Supplementary Table S1**.

### Tobacco grafting

Grafting between WT, NtCOP1OX, *NtCOP1^CR^* and NtHY5OX lines was performed according to method described by Rus et al. (2006). Seedlings grown on one-half-strength MS media for 7 days were used for grafting. Cuts between the scion and stock were made at an angle of 90° and were then placed in desired combinations. The grafted seedlings remained on the media plate for a period of 3 days to allow the formation of the graft union. Then, either only roots or whole grafted seedlings on the basis of scion used for example when *NtCOP1* were used as scion then whole seedlings and when *NtCOP1* were used as rootstock then only root portion was covered in darkness for 2 more weeks prior to sampling.

### Extraction and quantitative estimation of nicotine

Nicotine extraction was carried out with slight modifications (Khan *et al*., 2017). About 300 mg freeze-dried tissue samples were grinded to powder and extracted in 2.0 mL 75 % methanol. The extract was ultrasonicated for 60 min, and then centrifuged (14,000 rpm) for 10 min. The supernatant was concentrated to dry powder and then dissolved in 1mL HPLC grade methanol. Extracts were filtered through 0.2 mm filter (Millipore, USA) and subjected to HPLC analysis. Lab Solutions software (Shimadzu) was used for the quantification of nicotine through HPLC. Column used was Phenomenex Luna C18(2) column (250 mm x 4.6 mm x 5µ). Mobile phase consisted of water: methanol: 0.1 M buffer acetate (pH4.5): acetonitrile: acetic acid in the ratio 74: 3: 20: 2:1 (Solvent A) with pH adjusted to 4.2 with triethylamine. Solvent A was maintained at 100% throughout the run with a slight modification in the gradient of flow rate of 1ml/min in 0-15min, 1.0-1.5ml/min in 15-15.5min, 1.5ml/min in 15.5-24.5min, 1.5-1.0ml/min in 24.5-25min and 1ml/min in 25-30min. The samples were analysed by HPLC–PDA and chromatograms were recorded at 254nm (Pereira *et al*., 2001).

### Extraction and quantitative estimation of flavonols

Extraction of flavonols was carried out by grinding the plant material (300 mg) into the fine powder in liquid N_2_ and then extracted with 80% methanol overnight at room temperature with brief agitation. The extract was hydrolyzed with an equal amount of 6 N HCl at 70°C for 40 min followed by addition of an equal amount of methanol to prevent the precipitation of the aglycones (Jiang et al. 2015). Extracts were filtered through 0.2 mm filter (Millipore, USA) before HPLC. For non-hydrolyzed extracts, samples were extracted as described previously (Pandey *et al*., 2014). The mobile phase was a gradient prepared from 0.05% (v/v) ortho-phosphoric acid in HPLC-grade water (component A) and methanol (component B). Before use, the components were filtered through 0.45-mm nylon filters and de-aerated in an ultrasonic bath. The gradient from 25 to 50% B in 0–3 min, 50 to 80% B in 3–18 min, 80 to 25% B in 25 min and 25% B in 30 min was used for conditioning of the column with a flow rate of 1 ml/min. All the samples were analysed by HPLC–PDA with a Waters 1525 Binary HPLC Pump system comprising PDA detector (Niranjan *et al*., 2011).

### Extraction of total anthocyanin

For anthocyanin, 300 mg tissue were boiled for 3 min in extraction solution (propanol: HCl: H2O, 18:1:81), and incubated at room temperature in the dark for at least 2 h. Samples were centrifuged, and the absorbance of the supernatants was measured at 535 and 650 nm. Anthocyanin content was calculated as

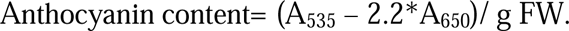

### Protein-protein interaction assay

To test the protein interaction by means of Yeast-two hybrid (H2H) assay, the bait vectors for expressing the self-AD deleted NtHY5 and NtCOP1 were constructed by cloning the corresponding sequences into pGBK-T7 vector (Takara Bio, San Jose, CA USA), and the prey vectors for expressing NtHY5 and NtCOP1 were constructed by cloning the full coding sequences into pGAD-T7 vector (Takara Bio, USA). The generated constructs were transformed into yeast (S. cerevisae) strain AH109 (Takara Bio, USA) as indicated pairs and cultured on SD/–Leu/–Trp medium at 30°C to select the desired colonies. The colonies were dropped onto SD/–Ade/– His/–Leu/–Trp medium supplemented with 20 mg/L X-aGal and cultured at 30°C for 3 d to test the protein interaction.

### Total protein extraction and western blot analysis

Roots (100 mg) of 20-day-old tobacco seedlings were frozen in liquid N2 and ground in extraction buffer (50 mM Tris (pH 7.5), 150 mM NaCl, 1 mM EDTA, 10% glycerol, 5 mM DTT, 1% protease inhibitor cocktail (Sigma-Aldrich), 0.1% Triton-X100). The homogenate was transferred to a tube and centrifuged at 20,000g for 12min at 4°C. The supernatant was transferred to a new tube, and an aliquot (5 µl) was used to estimate the protein concentration by Bradford assay75. Laemmli sample buffer (2X) was mixed with the protein sample and boiled at 96°C for 5 min. The protein samples (10µg and 50 µg) were separated on SDS–PAGE (12% and 20%, respectively) gel at 150V for 2h and transferred onto a PVDF membrane at 25V for 2h in transfer buffer (Tris–HCl (3.03g), glycine (14.4g), methanol (20%)) using a semi-dry transfer apparatus (Bio-Rad). After transfer, the membranes were incubated in the blocking solution (3% non-fat dry milk in Tris-buffered saline containing 0.05% Tween-20) for 1h at room temperature. The blots were incubated in primary antibodies diluted in TBS-T (1:10000 (v/v) for anti-actin, 1:1000 (v/v) for anti-HY5) overnight, followed by washing with wash buffer (Tris-buffered saline containing 0.05% Tween-20) thrice, 5 min each. The secondary antibody, conjugated with horseradish peroxidase (goat anti-rabbit IgG-HRP and goat anti-mouse IgG-HRP) diluted (1:10,000 times) in blocking buffer was incubated for 1h at room temperature, followed by washing with wash buffer three times, 5min each, followed by visualization using ChemiDoc (MP system, Bio-Rad) after incubating with luminol/enhancer and peroxide buffer (1:1 ratio). The western blot images were captured using Image Lab version 5.2.1 build 11 (Bio-Rad Laboratories). The commercial antibodies used in the analysis were Anti-Actin Antibody (A0480; Sigma-Aldrich), anti-HY5 (AS12 1867; Agrisera).

### Statistical analysis

Data are plotted as means±SE (n=3) in the figures. Data of flavonol and nicotine accumulation were evaluated for statistical significance using the Two-tailed Student’s t-test using Graphpad prism 5.01 software. Asterisks indicate significance levels with (* *p* < 0.1; ** *p* < 0.01; *** *p* < 0.001) as confidence intervals.

## RESULTS

### Identification and homolog analysis of NtCOP1

In our previous study, we delineated the involvement of NtHY5 in light-mediated nicotine biosynthesis in tobacco (Singh *et al*., 2023). Given the pivotal role of CONSTITUTIVE PHOTOMORPHOGENIC 1 (COP1) in modulating HY5 activity through ubiquitination and subsequent degradation (Bhatia *et al*., 2021), it emerges as a central regulator in light signaling cascades. To elucidate the functional significance of NtCOP1, we characterized the tobacco COP1, NtCOP1, and investigated its role in nicotine biosynthesis. Comparative analysis revealed that the NtCOP1 sequence shares approximately 80% similarity with Arabidopsis COP1 **(Supplementary Fig. S1A)**. Furthermore, phylogenetic examination based on amino acid sequences indicated a close relationship between NtCOP1 and COP1 proteins from diverse plant species across different families. Phylogenetic and pairwise distance analyses suggested that NtCOP1 exhibits a closer evolutionary relationship with CaHY5 and SlHY5 than with AtHY5 **(Supplemental Fig. S1B)**. Additionally, analysis from the conserved domain database indicated NtCOP1 as a member of the (RING)-finger E3 ubiquitin ligase family, similar to COP1 in other plant species. Noteworthy, NtCOP1 features an N-terminal ring finger motif and C-terminal 5 WD-40 domains **(Supplementary Fig. S2A)**. Sequence examination of *NtCOP1* revealed the presence of 13 exons and 12 introns, showcasing a structural similarity to *AtCOP1*, with variances in exon and intron sizes **(Supplementary Fig. S2B)**. Moreover, exploration of *NtCOP1* expression patterns across different tissues unveiled ubiquitous expression, with the highest expression observed in leaves and minimal expression detected in roots **(Supplementary Fig. S3)**.

### NtCOP1 is a functional ortholog of AtCOP1

To functionally characterize the identified NtCOP1, we generated homozygous transgenic lines in the Arabidopsis *cop1* mutant background (*atcop1*) for detailed analysis. Phenotypic complementation by NtCOP1 was evaluated based on growth parameters, including the number of leaves on the 10^th^ day of germination, hypocotyl length, and root length. Comparing the average hypocotyl lengths of 7-day-old seedlings from wild-type (COL0), *cop1* mutant, and NtCOP1;*cop1* complemented lines, we observed a significant recovery in hypocotyl length in the complemented lines **(Fig. 1A-B)**. Since light influences root growth in Arabidopsis, we measured the root length of 7-day-old light-grown seedlings from these genotypes, revealing a complete recovery of root length in the complemented lines to wild-type levels **(Fig. 1C)**. By the 10^th^ day of germination, the *cop1* mutant exhibited approximately 2 leaves, whereas the wild-type and complemented lines displayed 3 and 5 leaves, respectively, indicating a complementation of primary leaf development in the complemented lines **(Fig. 1D)**. Furthermore, mature (30-day-old) complemented plants, reached an average height of 8 cm, comparable to the 9 cm height of wild-type plants, while *cop1* mutant plants only reached a height of 3 cm **(Supplementary Fig. S4A and B)**. Additionally, we measured the hypocotyl length of 7-day-old dark-grown seedlings, which also showed a restoration in hypocotyl length after complementation **(Supplementary Fig. S4C and D)**. Results also suggested reduction in the anthocyanin levels upto the WT in complemented lines **(Fig. 1F)**. These results collectively suggest that NtCOP1 can complement the phenotypic defects of AtCOP1 in the *cop1* mutant.

**Fig. 1.**
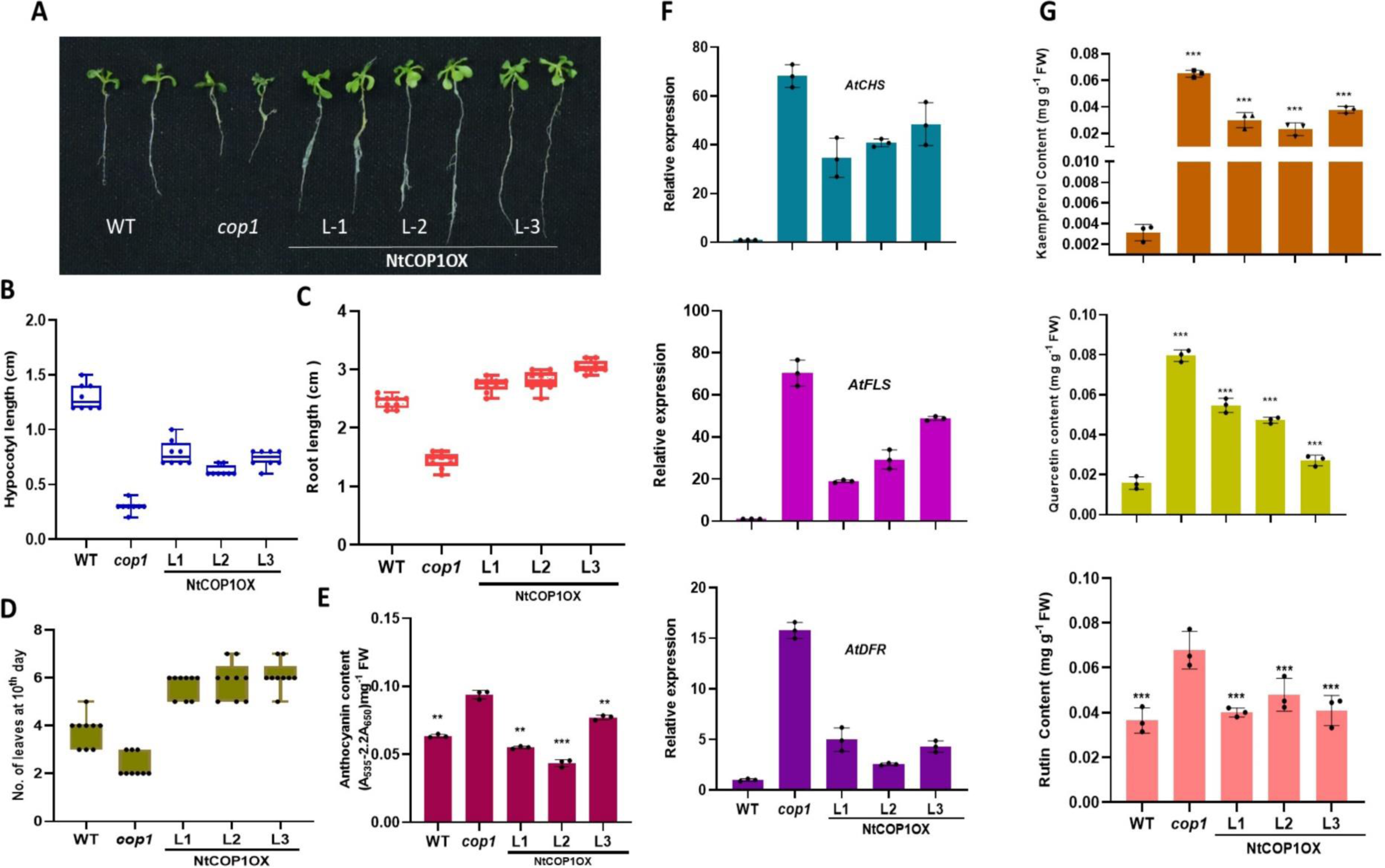
Complementation of Arabidopsis *cop1* mutant by NtCOP1. (A) Phenotype of 7-d-old light-grown WT (wild type; Col-0), *cop1* and NtCOP1OX;*cop1* seedlings (L1, L2 and L3 are three independent transgenic lines overexpressing NtCOP1 in *cop1* mutant background). (B) Hypocotyl length of 7-d-old white light grown WT, *cop1*, L1, L2 and L3 seedlings. (C) Root length of 7-d-old white light grown WT, *cop1*, L1, L2 and L3. (D) No. of leaves at 10^th^ day of germination of WT, *cop1*, L1, L2 and L3 seedlings. (E) Quantitative estimation of anthocyanin content in 7-d-old light grown WT, *cop1*, L1, L2 and L3 seedlings. (F) Relative expression of phenylpropanoid pathway genes (*AtCHS, AtFLS and AtDFR*) in 7-d-light grown WT, *cop1*, L1, L2 and L3 seedlings. (G) Quantitative estimation of flavonol (Kaempferol, Quercitin and Rutin) content in 7-d-old light grown WT, *hy5*, L1 and L2 seedlings through HPLC. Statistical analysis was performed using two-tailed Student’s t-test. Error bars represent SE of means (n=3). For hypocotyls, root length and no. of leaves at 10^th^ day of germination (n=10-12). Tubulin was used as endogenous control to normalize the relative expression levels. Error bars represent standard deviation. Asterisks indicate a significant difference, *P < 0.1, **P < 0.01, ***P < 0.001.

Subsequent gene expression and metabolite content analyses in the complemented lines revealed a recovery in the expression of key phenylpropanoid pathway genes, including *AtCHS, AtFLS*, and *AtDFR* **(Fig. 1F)**, as well as *AtPAL* **(Supplementary Fig. S4E)**, compared to the *cop1* mutant. Since flavonoids are known to be regulated by light signaling components, the heightened expression of flavonoid pathway genes prompted us to measure flavonol levels (Kaempferol, Quercitin, and Rutin). The results indicated elevated levels of flavonols in the complemented lines compared to the mutant lines **(Fig. 1G)**. These findings underscore the ability of NtCOP1 to not only rescue phenotypic defects in *cop1* mutant plants but also restore metabolite levels associated with light signaling regulation.

### NtCOP1 interact with NtHY5 to antagonistically regulate the nicotine biosynthesis

Previous studies have established that COP1 directly interacts with and modulates the activity of the transcription factor HY5, a key regulator of light-dependent gene expression and plant development (Ang *et al*., 1998; Holm *et al*., 2002; Han *et al*., 2019). However, the potential interaction between NtCOP1 and NtHY5 remains unexplored. Through bioinformatic analysis of the identified NtCOP1, we identified the presence of a WD40 domain, known for its role in binding to the VPE domain of AtHY5 for subsequent polyubiquitination. In a recent study, we proposed the existence of a VPE domain in NtHY5. To investigate the putative binding between NtHY5 and NtCOP1, we employed yeast two-hybrid analyses. The full-length coding sequences of NtHY5 and NtCOP1 were fused with a transcriptional activation domain (AD) in a prey vector and a DNA-binding domain (BD) in a bait vector, and vice versa. These constructs were co-expressed in yeast as prey–bait pairs (with AD-T7/BD-P53 serving as a positive control, AD-NtHY5/BD-NtCOP1 and AD-NtCOP1/BD-NtHY5 as binding proteins, and AD-T7/BD-Lam as a negative control). Successful yeast transformation was confirmed by growth on double dropout media (-His/-Leu), with yeast growth up to a 10^-4^ dilution in the negative control further confirming successful transformation **(Supplementary Fig. S5)**. Subsequently, to validate the binding between NtHY5 and NtCOP1, yeast cultures were grown on selective medium, SD/– Ade/–His/–Leu/–Trp (quadruple dropout media). The results revealed growth patterns similar to the positive control up to a 10^-4^ concentration, while no growth was observed in the negative control. These findings provide strong evidence supporting a binding interaction between NtCOP1 and NtHY5 **(Fig. 2A)**.

**Fig. 2.**
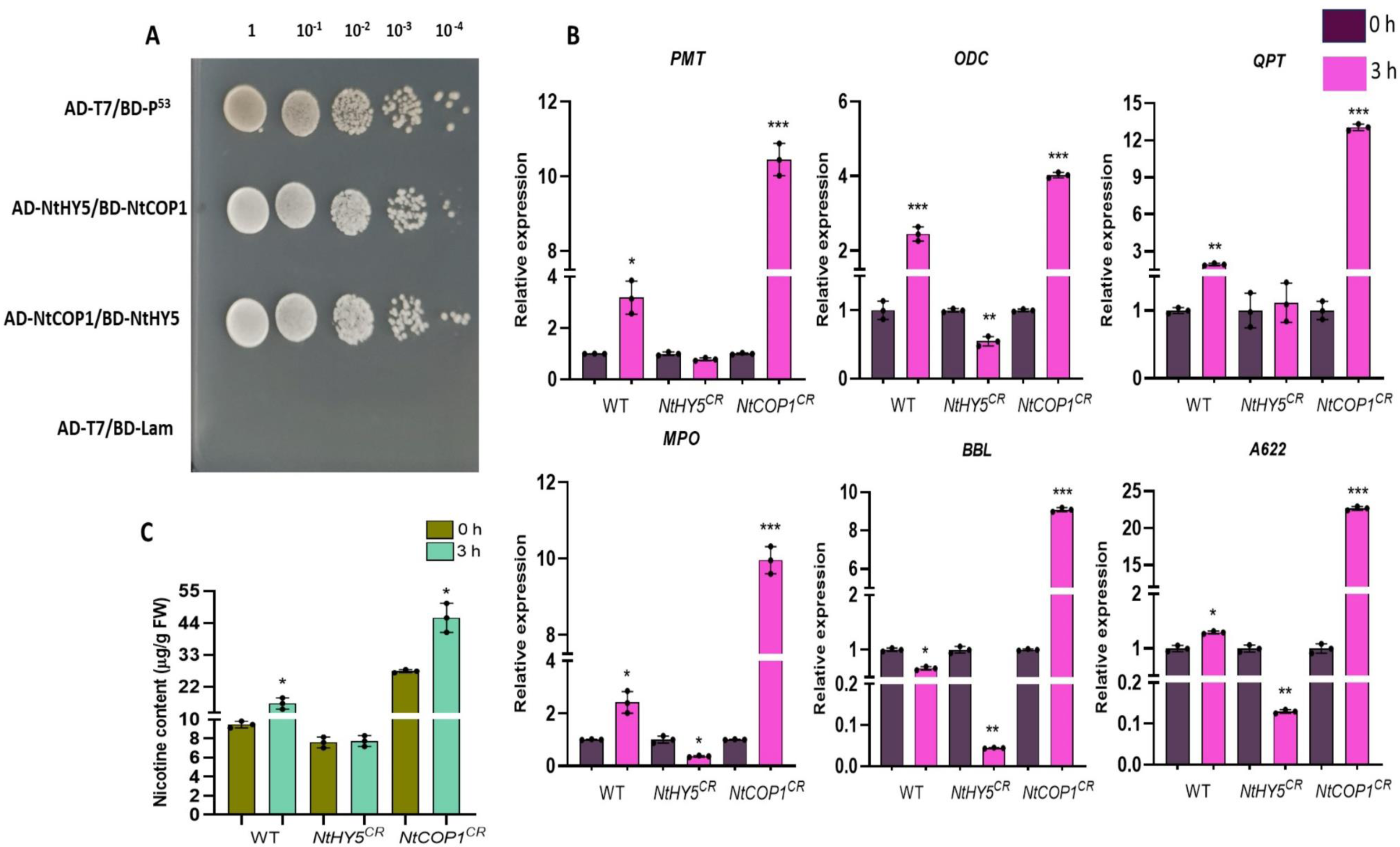
NtCOP1 interact with NtHY5 to antagonistically regulate the nicotine biosynthesis. (A) Yeast two hybrid (Y2H) analysis to show the *in vivo* iteraction between NtHY5 and NtCOP1. (B) Relative expression analysis of *NtPMT, NtODC, NtQPT, NtMPO, NtBbl* and *NtA622* in WT, *NtHY5^CR^* and *NtCOP1^CR^* seedlings grown for 10 days in dark followed by an exposure to light for 3hr. (C) Quantitative estimation of nicotine content through HPLC in WT, *NtHY5^CR^* and *NtCOP1^CR^* seedlings grown for 10 days in dark followed by an exposure to light for 3hr. Tubulin was used as endogenous control to normalize the relative expression levels. Error bars represent standard deviation. Asterisks indicate a significant difference, *P < 0.1, **P < 0.01, ***P < 0.001.

In our recent study (Singh *et al*., 2023), we found that NtHY5 positively regulates nicotine biosynthesis, as evidenced by lower nicotine levels in *NtHY5^CR^* plants. Building on this, in the current study, we developed CRISPR/Cas9-based NtCOP1 mutant lines in tobacco **(Supplementary Fig. S6A-D)**. Relative expression analysis suggested the lower expression of *NtCOP1* in developed *NtCOP1^CR^* lines in comparison to WT **(Supplementary Fig. S6E)**. Next, to analyze the effect of light to dark transition, we grew WT, *NtHY5^CR^*, and *NtCOP1^CR^*seedlings for 10 days in light and then transferred them to darkness for 3 hours. Real-time expression analysis of these seedlings revealed a higher induction in the expression of nicotine pathway genes, including *NtPMT, NtODC, NtQPT, NtMPO, NtBbl,* and *NtA622*, from dark to light in *NtCOP1^CR^*seedlings. In contrast, *NtHY5^CR^* seedlings showed no induction in the gene expression **(Fig. 2B)**. Further, estimation of nicotine content after dark-to-light induction further supported these findings, indicating a higher induction in nicotine content in *NtCOP1^CR^*seedlings, while *NtHY5^CR^* seedlings exhibited significantly less induction compared to WT **(Fig. 2C)**. Analysis also suggested that *NtHY5^CR^* seedlings accumulate more nicotine compared to wild type seedlings, in the dark as well as light conditions.

### NtCOP1 mediates the light-dependent regulation of nicotine and flavonoid biosynthesis

To functionally characterize tobacco’s COP1, we generated NtCOP1 overexpression lines (NtCOP1OX) alongside NtHY5 CRISPR-edited lines (*NtHY5^CR^*). The NtCOP1OX lines demonstrated significantly enhanced NtCOP1 expression levels compared to the WT **(Supplementary Fig. S6F)**. To analyse phenotypic and molecular differences between overexpression and edited lines, NtCOP1OX, *NtCOP1^CR^*, and WT seedlings were cultivated for 10 days under light conditions. Our findings revealed that both hypocotyl and root lengths were significantly shorter in *NtHY5^CR^* seedlings in comparison to both NtCOP1OX and WT counterparts **(Fig. 3A-C)**.

**Fig. 3.**
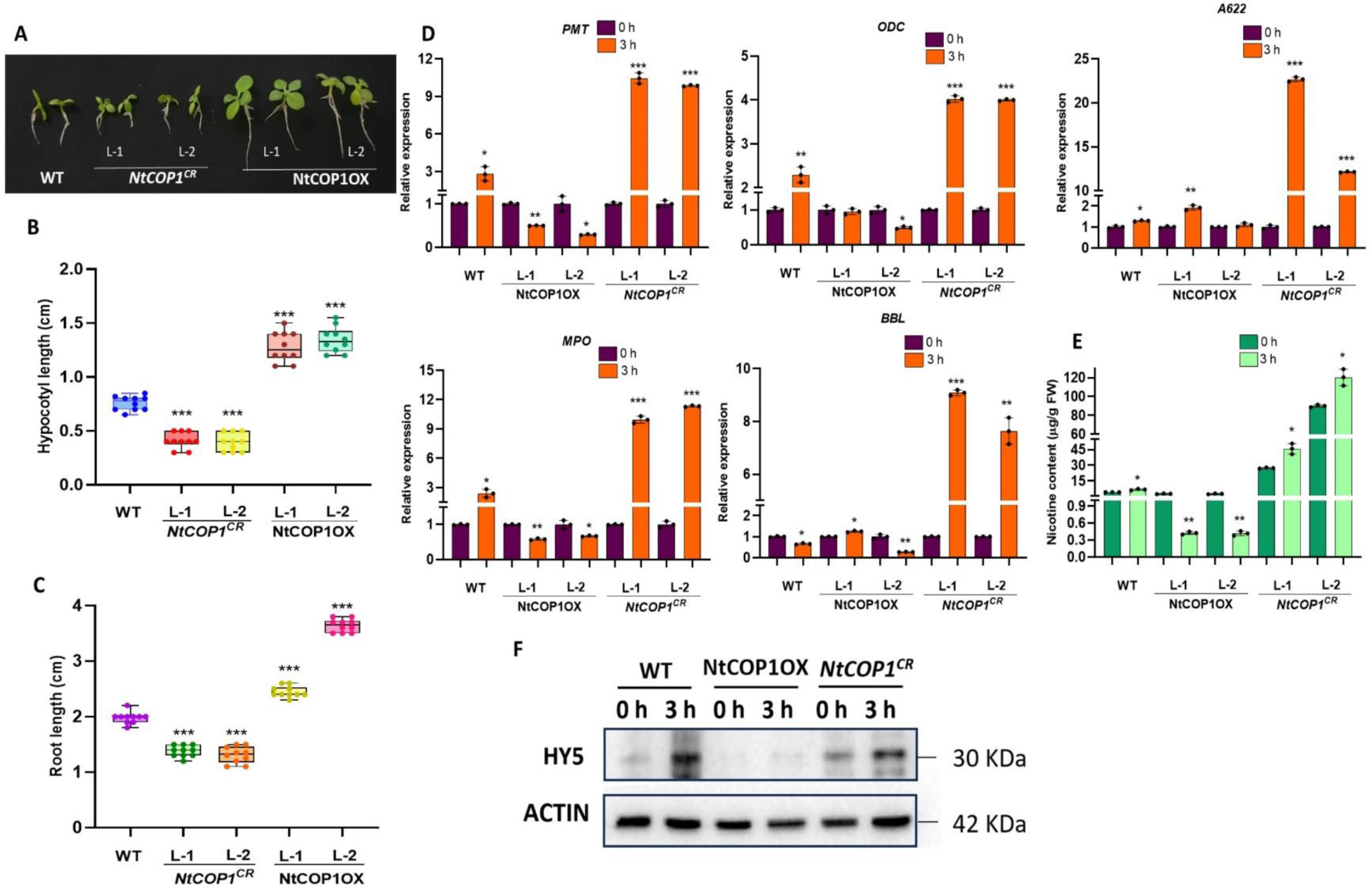
Overexpression of NtCOP1 and CRISPR-Cas9-mediated knockout of *NtCOP1* modulates gene expression and accumulation of metabolite in tobacco seedlings. (A)Visual representation of the 10-d-old white light grown seedlings. (B) Hypocotyl length of 10-d-old white light grown WT, *NtCOP1^CR^* and NtCOP1OX seedlings. (C) Root length of 10-d-old white light grown WT, *NtCOP1^CR^* and NtCOP1OX seedlings. (D) Relative transcript abundance of nicotine pathway genes (*NtPMT, NtODC, NtQPT, NtMPO, NtBBL and NtA622*) in WT, *NtCOP1^CR^* and NtCOP1OX seedlings grown for 10 days in dark followed by an exposure to light for 3hr. (E) Quantification of nicotine content through HPLC in WT, *NtCOP1^CR^* and NtCOP1OX seedlings grown for 10 days in dark followed by an exposure to light for 3hr. (F) NtHY5 levels in WT, *NtCOP1^CR^* and NtCOP1OX seedlings grown for 10 days in dark followed by an exposure to light for 3hr Statistical analysis was performed using two-tailed Student’s t-test. Error bars represent SE of means (n=3). For hypocotyls and root length (n=10-12). Tubulin was used as endogenous control to normalize the relative expression levels. Error bars represent standard deviation. Asterisks indicate a significant difference, *P < 0.1, **P < 0.01, ***P < 0.001.

To assess the expression of nicotine and flavonoid pathway genes, NtCOP1OX, *NtCOP1^CR^*, and WT seedlings were grown for 10 days under continuous dark conditions, followed by a 3-hour light exposure. Our findings revealed a higher induction in the expression of nicotine pathway genes (*NtPMT, NtODC, NtMPO, NtBBL, and NtA622*) in *NtCOP1^CR^*lines, whereas NtCOP1OX lines exhibited lower expression levels compared to WT after 3 hours of light exposure **(Fig. 3D)**. Moreover, analysis of flavonoid pathway genes (*NtCHS, NtFLS, and NtDFR*) displayed a similar trend, with higher induction observed in *NtCOP1^CR^* lines and reduced expression in NtCOP1OX lines compared to WT **(Supplementary Fig. S7A).** These expression patterns led us to estimate nicotine and flavonol contents. The analysis suggested a significant increase in both nicotine **(Fig. 3E)** and flavonol accumulation **(Supplementary Fig. S7B)** in *NtCOP1^CR^*seedlings, while NtCOP1OX lines exhibited reduced accumulation levels compared to WT. We also measured levels of NtHY5 under these conditions. Our western blot analysis suggests an increase in NtHY5 levels after exposure to light in WT and *NtCOP1^CR^* plants. However, no such increase was observed in NtCOP1Ox lines **(Figure 3F)**.

### Effect of NtCOP1 on nicotine accumulation in mature tobacco plants

To assess the impact of NtCOP1 on mature plants, 3-month-old NtCOP1OX, *NtCOP1^CR^*, and WT plants, grown under normal conditions, were subjected to complete darkness for 24 hours. Subsequently, root samples were collected for the analysis of nicotine pathway gene expression. The expression analysis revealed higher induction of gene expression in *NtCOP1^CR^* plants, while NtCOP1OX plants exhibited lower induction compared to WT **(Figure 4A-F)**. Nicotine accumulation in the leaves of these plants was also measured, indicating significantly higher induction in *NtCOP1^CR^* plants compared to NtCOP1OX and WT **(Figure 4G)**. Furthermore, examination of anthocyanin content in flower petals demonstrated higher accumulation in *NtCOP1^CR^* flowers than in NtCOP1OX, relative to WT **(Supplementary Fig. S8 A and B)**. Phenotypic differences were observed, with NtCOP1OX plants being taller and *NtCOP1^CR^* plants smaller compared to WT **(Fig. 4H)**. These results suggest a NtCOP1 mediated light-dependent regulation of nicotine biosynthesis and plant phenotype.

**Fig. 4.**
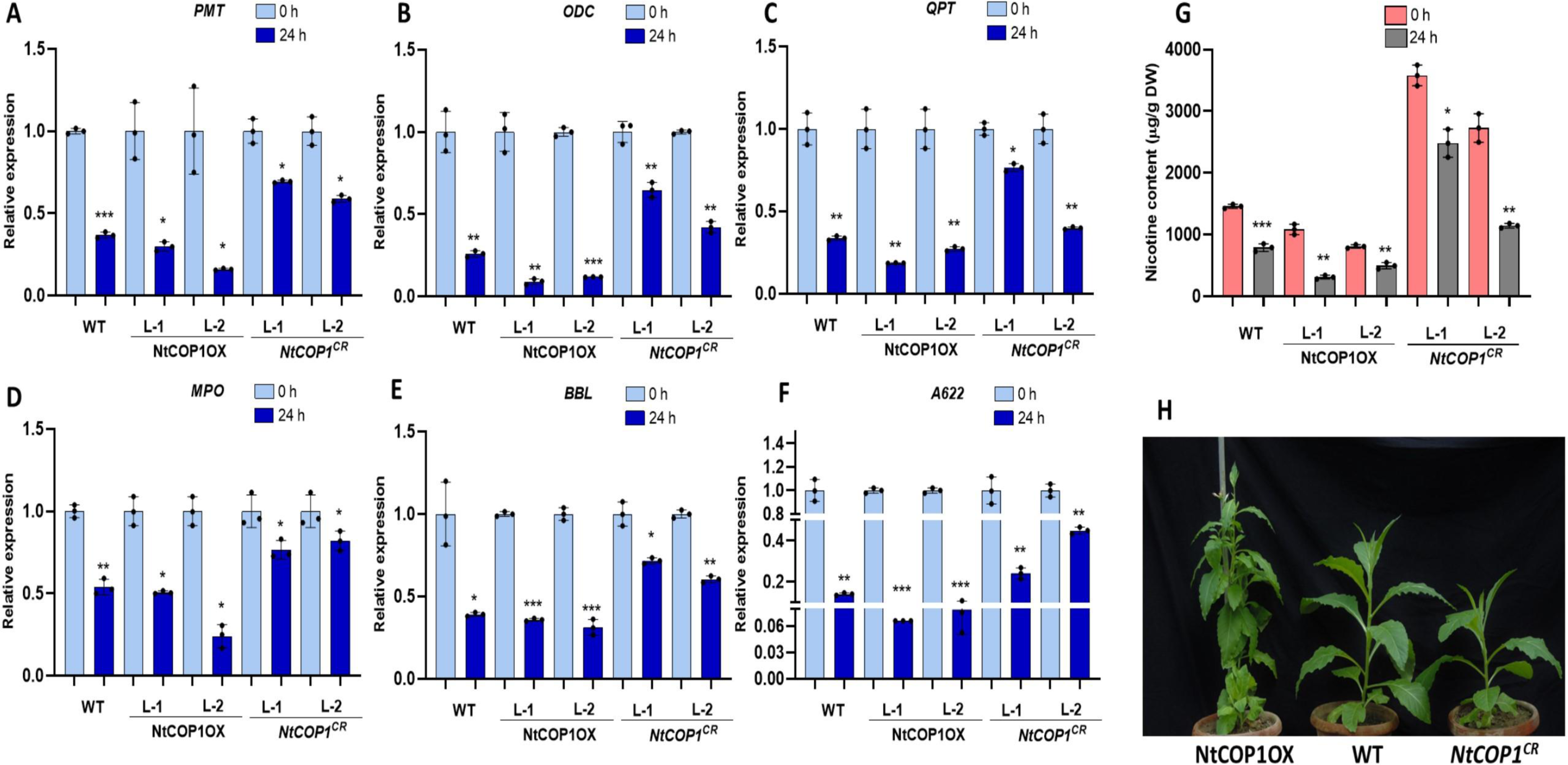
NtCOP1 modulates nicotine levels in mature tobacco transgenic lines. (A-F) Relative transcript abundance of nicotine pathway genes *NtPMT, NtODC, NtQPT, NtMPO, NtBbl* and *NtA622* respectively in roots of WT, NtCOP1OX and *NtCOP1^CR^* 3-month-old light grown tobacco plants transferred to dark for 24h. (G) Quantitative estimation of nicotine content in leaves of 3-month-old WT, NtCOP1OX and *NtCOP1^CR^* light grown tobacco plants transferred to dark for 24h through HPLC. (H) Visual phenotypic difference in the growth of NtCOP1OX, WT and *NtCOP1^CR^* 3-month-old light grown tobacco plants. Statistical analysis was performed using two-tailed Student’s t-test. Error bars represent SE of means (n=3). Tubulin was used as endogenous control to normalize the relative expression levels. Error bars represent standard deviation. Asterisks indicate a significant difference, *P < 0.1, **P < 0.01, ***P < 0.001.

### Shoot NtCOP1 regulates nicotine biosynthesis in root via regulating NtHY5 protein

Our earlier findings via Y2H indicated the binding of NtCOP1 to NtHY5 (**Fig. 2A**). Given that Arabidopsis COP1 is a constitutive repressor of photomorphogenesis known to interact with photomorphogenesis-promoting factors like HY5, facilitating their proteasome-mediated degradation (Saijo *et al*., 2003), we aimed to establish the correlation between NtHY5, NtCOP1, and nicotine biosynthesis in tobacco. To achieve this, we conducted grafting and western blot experiments. In the grafting experiment, shoots (scion) and roots (rootstocks) of 7-day-old WT, NtHY5OX, NtCOP1OX, and *NtCOP1^CR^* seedlings were combined in various configurations **(Supplementary Fig. S9A)**. Combinations included NtHY5OX/*NtCOP1^CR^*, NtHY5OX/NtCOP1OX, NtCOP1OX/NtHY5OX, NtCOP1OX/*NtHY5^CR^*, NtCOP1OX/WT, *NtCOP1^CR^*/WT, and WT/WT. After three days of grafting, complete seedlings were covered to prevent light exposure. In combinations where NtCOP1OX or *NtCOP1^CR^* seedlings were used as rootstocks, only roots were covered **(Supplementary Fig. S9B).** In combinations where NtCOP1OX or *NtCOP1^CR^* seedlings were used as scion, the entire seedlings were covered **(Supplementary Fig. S9C)**. After two weeks, roots and leaves were sampled for expression analysis and nicotine content measurement, respectively. Two control scenarios were considered: one with NtHY5OX/*NtCOP1^CR^*as control and another with WT/WT as control.

Expression analysis of nicotine pathway genes in various grafting combinations revealed significantly higher expression when NtHY5OX served as a scion and *NtCOP1^CR^* as rootstock (NtHY5OX/*NtCOP1^CR^*), and the lowest expression when NtCOP1OX was the scion with *NtHY5^CR^*as the rootstock (NtCOP1OX/*NtHY5^CR^*) **(Fig. 5A-F)**. Nicotine accumulation mirrored this pattern, with the highest levels observed in NtHY5OX/*NtCOP1^CR^* and the lowest in *NtHY5^CR^/NtHY5^CR^*grafted union, followed by NtCOP1OX/*NtHY5^CR^* **(Fig. 5G)**. When using WT/WT as a control, NtCOP1OX/WT exhibited lower expression of nicotine pathway genes compared to NtHY5OX/WT **(Supplementary Fig. S10A-F)**. Consistent with this expression pattern, nicotine accumulation was also lower in NtCOP1OX/WT compared to *NtCOP1^CR^*/WT **(Supplementary Fig. S10G)**. These findings suggest that NtCOP1 in the shoot degrades shoot NtHY5 in the dark, preventing it from reaching the root, activating local HY5, and consequently reducing nicotine biosynthesis.

**Fig. 5.**
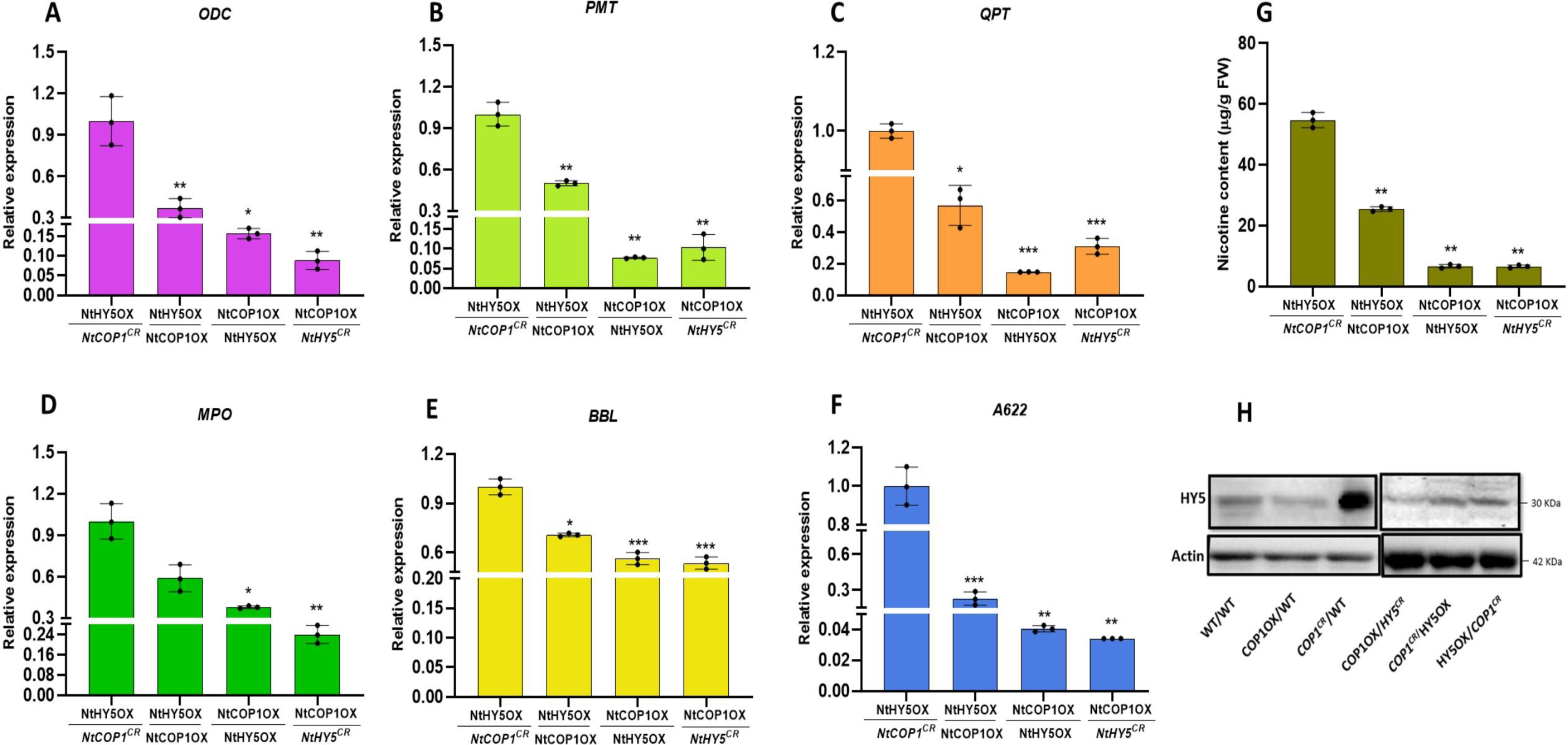
NtCOP1 idirectly regulates nicotine levels in tobacco by directly regulating the concentration of NtHY5 in root. (A-F) Relative expression of nicotine pathway genes (*NtODC, NtPMT, NtQPT, NtMPO, NtBBL and NtA622* respectively) in roots of grafted NtCOP1OX, *NtCOP1^CR^*, NtHY5OX and *NtHY5^CR^* seedlings in several combinations keeping NtHY5OX*/NtCOP1^CR^* as control. (G) Quantitative estimation of nicotine content in grafted NtCOP1OX, *NtCOP1^CR^*, NtHY5OX and *NtHY5^CR^* seedlings. Epicotyl tissue (300 mg fresh weight) was used for the analysis. (H) Western blot analysis to show the concentration of NtHY5 protein in roots of grafted WT, NtCOP1OX, *NtCOP1^CR^*, NtHY5OX and *NtHY5^CR^* seedlings in different combinations. After three days of grafting, complete seedlings were covered to prevent light exposure. In combinations where NtCOP1OX or *NtCOP1^CR^* seedlings were used as rootstocks, only roots were covered. In combinations where NtCOP1OX or *NtCOP1^CR^* seedlings were used as scion, the entire seedlings were covered. Roots and hypocotyls were used for all the expression analysis and nicotine quantification, respectively. TUBULIN was used as the endogenous control to normalize the relative expression levels. The statistical analysis was performed using two-tailed Student’s t-tests. The data are plotted as meansLJ±LJSD (n= 3). The error bars represent standard deviations. The asterisks indicate significant difference, *P<LJ0.1; **P<LJ0.01; ***P<LJ0.001.

To ascertain whether NtCOP1 plays a direct or indirect role in nicotine regulation, we conducted a western blot analysis. Crude protein isolated from grafted seedlings showed higher levels of NtHY5 protein in *NtCOP1^CR^*/WT, followed by WT/WT, and the least in NtCOP1OX/WT. Similarly, the highest protein accumulation occurred in NtHY5OX/*NtCOP1^CR^*, followed by *NtCOP1^CR^*/HY5OX, and the least in NtCOP1OX/*NtHY5^CR^* (**Fig. 5H)**. These results suggest that in NtCOP1OX seedlings as scion, NtCOP1 degrades NtHY5 in the leaf, preventing it from moving to the root and activating root-local HY5, resulting in lower protein levels. In contrast, when *NtCOP1^CR^* is the scion, NtCOP1 does not degrade NtHY5, allowing it to move down to the root and accumulate, leading to higher levels of NtHY5 protein.

## DISCUSSION

Nicotine, the principal addictive component in tobacco products, exerts widespread biological effects on various bodily systems, including cardiovascular, respiratory, renal, and reproductive systems. Numerous studies have consistently demonstrated its carcinogenic potential (Heeschen *et al*., 2001; Nakada *et al*., 2012). Reducing nicotine levels is a crucial objective in tobacco research using molecular science. Nicotine synthesis occurs in root cortical cells and requires transportation to the leaves, where it accumulates and serves as an antiherbivory defense. The dynamic movement of nicotine through biological membranes involves tonoplast-localized multidrug and toxic compound extrusion transporters (MATEs) such as MATE1, MATE2, and jasmonate-inducible alkaloid transporter 1 (JAT1) (Morita *et al*., 2009; Shoji *et al*., 2009), along with plasma membrane-localized purine permease family member nicotine uptake permease 1 (NUP1). Recent studies have extensively explored transcriptional regulators of nicotine biosynthesis, identifying distinct families of transcription factors including APETALA2 (AP2)/ethylene response factor (ERF), NtORC1/ERF221, NtJAP1/ERF10, MYC2-like basic helix-loop-helix (bHLH) such as AtMYC2a/b, NbbHLH1, and NbbHLH2 families, as well as MYB (NtMYB305a) (Song *et al*., 2011; Shoji *et al*., 2011a; Sears *et al*., 2014; Liu *et al*., 2019; Bian *et al*., 2021).

Earlier studies in Arabidopsis and other plant species have established HY5 as a master regulator of photomorphogenesis and a crucial signaling component in light response, promoting photomorphogenesis across a broad spectrum of wavelengths (Ulm *et al*., 2004; Lee *et al*., 2007; Nawkar *et al*., 2017). However, our recent study (Singh *et al*., 2023) focuses on the identification and functional characterization of tobacco HY5. In this study, we revealed that NtHY5 functions as a shoot-root mobile signal, orchestrating the light-regulated synthesis of nicotine in the root. This regulatory mechanism involves NtHY5 interacting with the G-BOXES present within the promoters of nicotine pathway genes, specifically *NtPMT, NtQPT,* and *NtODC*.

Previous studies have established COP1 and HY5 as antagonistic nuclear regulators, with COP1 directly interacting with the bZIP transcription factor HY5, a positive regulator of photomorphogenesis, and promoting its proteasome-mediated degradation (Ang *et al*., 1998; Holm *et al*., 2002). This prompted us to hypothesize the role of NtCOP1 in nicotine biosynthesis, antagonistic to NtHY5. In the current study, we identified tobacco COP1 and explored its regulatory role in nicotine production. While various COP1 orthologs, including those from Physcomitrella and rice, have been previously identified and functionally conserved during evolution (Wu *et al*., 2019; Ranjan *et al*., 2014), this study marks the first identification of NtCOP1. Sequence analysis revealed over 90% similarity between Arabidopsis and tobacco COP1, with a conserved domain in NtCOP1. The only difference observed was the presence of five WD40 domains in NtCOP1, as opposed to the seven in Arabidopsis COP1. This suggests functional conservation of protein sequences between tobacco and Arabidopsis during evolution. Complementation studies were conducted to assess whether NtCOP1 could fully complement the Arabidopsis *cop1* mutant. The study demonstrated partial complementation by NtCOP1 for phenotypes such as hypocotyl length in light and dark. However, apical hook formation, plant height, relative expression of *AtCHS, AtFLS, and AtDFR*, and metabolite content (Flavonols and Anthocyanin) were not fully restored due to partial complementation **(Fig. 1; Supplementary Fig. S3**). Similar results were reported by Tsuge et al. (2006), indicating that rice COP1 could complement the Arabidopsis cop1 mutant in processes like greening, apical hook formation, and hypocotyl length. These findings suggest that NtCOP1 maintains conserved functions across species, not only at the phenotypic level but also at the metabolite level, such as flavonoid content.

To assess the individual impact of NtHY5 and NtCOP1 mutations on gene expression and nicotine content in tobacco seedlings, *NtCOP1^CR^*(developed in this study using CRISPR/Cas9 technology) and *NtCOP1^CR^*(previously developed in our research) seedlings were cultivated in light for 10 days, followed by 3 hours of darkness. Results indicated higher expression and nicotine levels in NtCOP1CR seedlings compared to *NtHY5^CR^*. These findings suggest an interaction between NtCOP1 and NtHY5. In NtCOP1OX lines, NtCOP1 degrades NtHY5, preventing its binding to the promoter of nicotine pathway genes and resulting in reduced nicotine content. Conversely, in *NtCOP1^CR^* seedlings, the absence of NtCOP1 allows NtHY5 to bind to the promoter, leading to increased nicotine content. To delve into the molecular mechanism, a Yeast Two-Hybrid (Y-2-H) assay was conducted (**Fig. 2B**). The growth of yeast on quadruple dropout media strongly indicated the interaction between NtHY5 and NtCOP1, akin to the interaction between AtHY5 and AtCOP1 (Ang *et al*., 1998; Holm *et al*., 2002).

To go deeper into the function of NtCOP1, we developed NtCOP1 overexpression lines in tobacco. After cultivating the seedlings for 10 days in light, we observed phenotypic differences in hypocotyl and root length. Notably, NtCOP1OX seedlings exhibited root lengths almost twice as long as *NtCOP1^CR^*seedlings, suggesting the role of NtCOP1 in seedling growth. This aligns with previous reports indicating that COP1 regulates various development-related proteins, including light-responsive proteins like HY5 (Kim *et al*., 2017). We further explored the variations in the relative expression of nicotine pathway genes and nicotine content. After subjecting the 10-day light-grown seedlings to 3 hours of darkness, the results revealed approximately 10 times higher induction in expression and a significant increase in nicotine content in *NtCOP1^CR^* seedlings compared to NtCOP1OX and WT (**Fig. 3D and E**). These findings imply that in the dark, NtCOP1 translocates to the nucleus, degrading NtHY5 in NtCOP1OX lines, leading to low nicotine accumulation. In *NtCOP1^CR^* seedlings, the mutated COP1 interferes with the interaction between normal COP1 and SPA1, causing incomplete COP1-mediated ubiquitination and degradation of HY5. Consequently, during the dark-to-light transition, NtHY5 levels increase, enhancing nicotine content by binding to the promoter of nicotine pathway genes. Given previous studies indicating the role of COP1 in anthocyanin and flavonoid regulation, we examined the expression of major flavonoid pathway genes (NtCHS*, NtFLS,* and *NtDFR*) and flavonoid content (flavonols and anthocyanin). The results revealed that NtCOP1 negatively regulates flavonols and anthocyanin production in tobacco (**Supplementary Fig. S7 and S8**). To assess NtCOP1’s impact on mature plants, we subjected 3-month-old light-grown plants to 24 hours of darkness. The results indicated that during the light-to-dark transition, *NtCOP1^CR^* plants showed less reduction in the expression of nicotine pathway genes and nicotine content compared to NtCOP1OX and WT, suggesting a negative role of NtCOP1 in nicotine accumulation in tobacco leaves (Figure 4A-F and 4G). Additionally, a significant difference in the height of mature plants was observed, with NtCOP1OX plants being taller than *NtCOP1^CR^*, indicating that NtCOP1 degrades NtHY5 and other developmental-related proteins, ultimately influencing plant growth. Previous studies have suggested that COP1 degrades proteins like HY5, HYH, LZF1/STH3 to regulate plant growth (Holm *et al*., 2002; Datta *et al*., 2008; Srivastava *et al*., 2015).

To elucidate the molecular mechanism by which NtCOP1 influences nicotine synthesis in tobacco roots known for nicotine production (Hayashi *et al*., 2020) we examined how shoot NtCOP1 regulates nicotine production in roots through grafting experiments. In dark conditions, NtCOP1OX lines exhibited degradation of NtHY5, rendering it unavailable to move from the shoot to the root. Consequently, root-localized NtHY5 failed to bind to the promoter of nicotine pathway genes, resulting in decreased nicotine synthesis. Previous studies have indicated that HY5 moves from the shoot to the root and modulates nicotine synthesis in the root (Chen *et al*., 2016; Singh *et al*., 2023). Protein extraction from grafted seedling roots revealed lower NtHY5 protein levels when NtCOP1OX served as the scion, and higher levels when *NtCOP1^CR^* served as the scion. This aligned with our previous finding that lower NtHY5 protein levels correlated with reduced nicotine accumulation in NtCOP1OX/ *NtCOP1^CR^* and NtCOP1OX/WT, and higher accumulation in *NtCOP1^CR^* /NtCOP1OX and *NtCOP1^CR^* /WT grafted seedlings (**Fig. 5** and **Supplementary Fig. S10**). Thus, the study indicates that NtCOP1 indirectly regulates nicotine biosynthesis in roots by degrading NtHY5 in leaves. In the case of mutated COP1, where a truncated NtCOP1 lacks the binding-responsible WD40 domain, NtHY5 can move to the root in higher concentrations, bind to the promoter of nicotine pathway genes, and upregulate nicotine biosynthesis. The NtCOP1 overexpression plants developed, in this study; with low nicotine accumulation in leaves could help people to overcome their nicotine addiction and the risk of death from tobacco use.

In conclusion, our proposed model **(Figure 6)** illustrates the regulatory mechanism involving COP1, an E3 ubiquitin ligase, and NtHY5. COP1 binds to NtHY5 and mediates its degradation, consequently inhibiting the movement of NtHY5 from shoot to root. As a result, NtHY5 becomes unavailable to bind to the promoters of nicotine pathway genes in the root, leading to a suppression of nicotine synthesis. Conversely, when NtCOP1 is mutated, it fails to degrade NtHY5 in the shoot, allowing NtHY5 to move freely from shoot to root and activate the promoters of nicotine pathway genes, thus promoting nicotine biosynthesis. Additionally, our findings indicate that NtCOP1 similarly regulates flavonoid biosynthesis in tobacco leaves by degrading NtHY5. This comprehensive model highlights the intricate role of COP1 in modulating both nicotine and flavonoid biosynthesis pathways through its interaction with NtHY5.

**Fig. 6.**
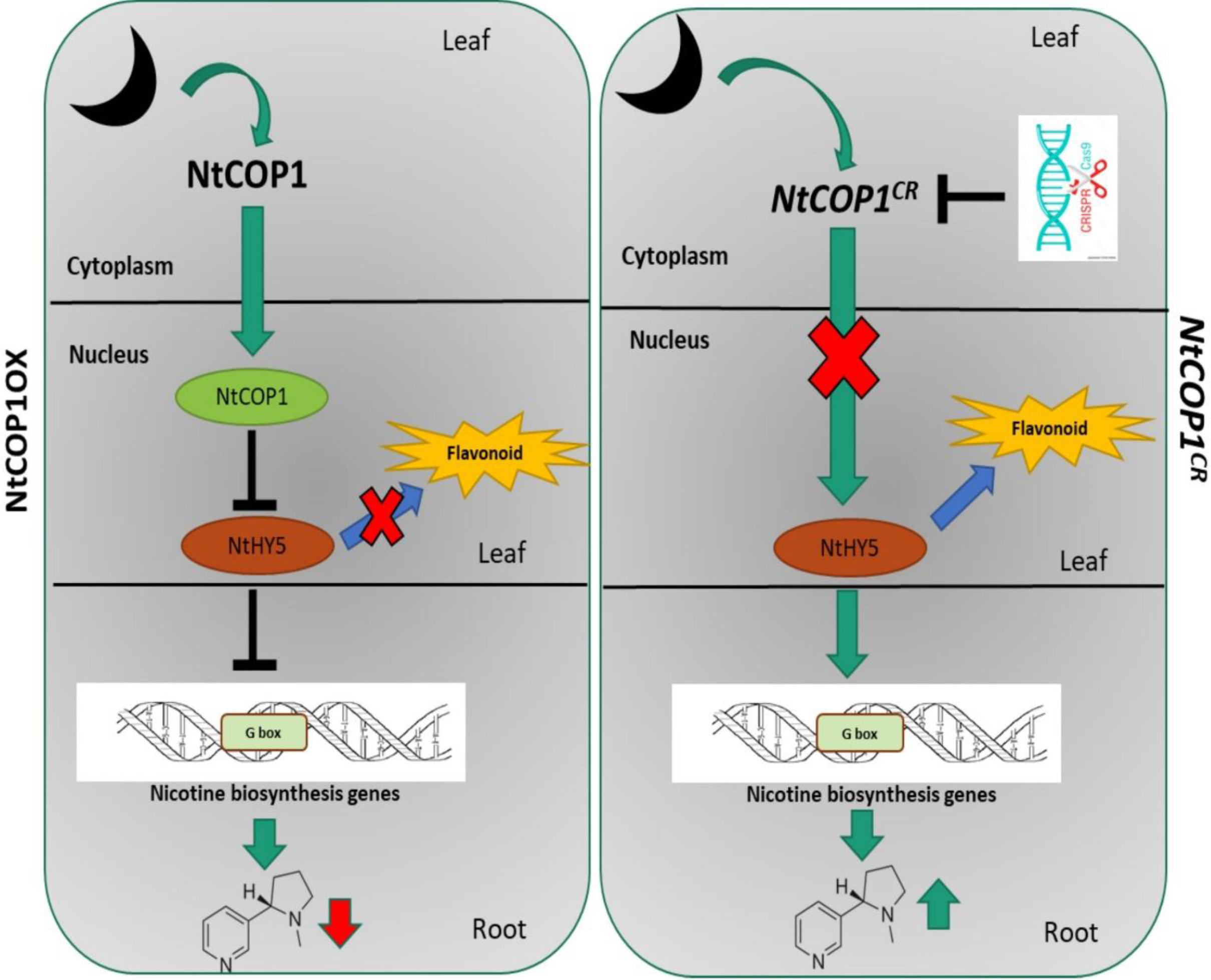
Proposed working model illustrating the involvement of NtCOP1 in nicotine and flavonol accumulation in tobacco. The model delineates that in darkness, NtCOP1 binds and degrades NtHY5 via the proteasomal machinery, rendering NtHY5 unavailable for translocation from shoot to root. Consequently, NtHY5 is unable to interact with the promoters of nicotine pathway genes, leading to a reduction in nicotine synthesis. Conversely, mutation in NtCOP1 renders it non-functional, resulting in the stabilization of NtHY5 even in dark conditions. This allows NtHY5 to freely translocate from shoot to root and interact with the promoters of nicotine pathway genes, thereby enhancing nicotine biosynthesis. Similarly, NtCOP1 governs flavonoid biosynthesis in tobacco leaves. Arrowheads and tees represent positive and negative regulation, respectively.

## Data availability

All data generated or analyzed during this study are included in this published article (and its supplementary information files).

## Acknowledgement

P.K.T. acknowledges CSIR and DBT, New Delhi, for financial support in the form of projects on pathway engineering and genome editing. P.K.T. also acknowledges Science and Engineering Research Board, New Delhi for JC Bose National Fellowship (JCB/2021/000036). DS acknowledge University Grant Commission, New Delhi for Senior and Junior Research Fellowship respectively. Authors acknowledge Dr. Manju Singh, Chemical Central Facility, CSIR-CIMAP for HPLC analysis.

## Author Contributions

P.K.T. designed and supervised this study. D.S. designed and per formed most of the experiments; NS and SD provided help in conducting various experiments; P.K.T. analyzed the data; D.S., P.K.T. wrote the article; all authors read, contributed, and approved the article.

## Conflict of interest

All authors contributing to this work have no conflicts of interest to disclose.

## Funding statement

P.K.T. acknowledges CSIR and DBT, New Delhi, for financial support in the form of projects on pathway engineering and genome editing. P.K.T. also acknowledges Science and Engineering Research Board, New Delhi for JC Bose National Fellowship (JCB/2021/000036).

## Supplementary data

Figure S1. Multiple Sequence alignment and phylogenetic tree of NtCOP1 gene.

Figure S2. Sequence analysis of NtCOP1 gene.

Figure S3. Tissue specific normalized expression of NtCOP1 gene.

Figure S4. Complementation of *Arabidopsis cop1* mutant by NtCOP1.

Figure S5. Yeast two hybrid analysis (Y2H) to show binding between NtHY5 and NtCOP1.

Figure S6. Development of CRISPR/Cas9 mediated NtCOP1 edited plants.

Figure S7. Modulation of flavonol biosynthesis in tobacco by NtCOP1.

Figure S8. Modulation of anthocyanin biosynthesis in tobacco by NtCOP1.

Figure. S9. Phenotype of the grafted seedlings.

Figure S10. Modulation in the expression of nicotine pathway genes and nicotine content in grafted seedlings.

Table S1. List of primers used for development of constructs and expression analysis in this study.

## Notes

### Competing Interest Statement

The authors have declared no competing interest.

